# Fertility reversibly modulates *C. elegans* behavior via gonad-nervous system signaling

**DOI:** 10.64898/2026.07.23.740032

**Authors:** Isobella Veitch, Emily A. Bayer

**Affiliations:** Department of Biology and Center for Neural Science, New York University, New York, NY, USA 10003; Biozentrum, University of Basel, 4056 Basel, Switzerland

## Abstract

While the external cues that generate behavioral responses have been extensively studied, behavioral changes in response to internal stimuli are much less understood. *C. elegans* hermaphrodites are self-fertile until they exhaust their supply of self-sperm, after which time they can only produce more progeny via mating with a male animal. Fertile hermaphrodites are less likely to be mated by males than sperm-exhausted hermaphrodites. We report that sperm-dependent hermaphrodite escape is facilitated by an aversive response to physical contact (touch stimulus) by males, specifically male turns around the hermaphrodite nose and male contact at the vulva. Using germline masculinizing and feminizing mutants, we found that sperm are both necessary and sufficient to induce contact-dependent mating evasion. Surprisingly, loss of the entire somatic gonad resulted in ectopic evasion behavior, via an oppositional monoaminergic signal. Hermaphrodites lacking the HSN neuron are constitutively receptive to male mating touch both at the vulva and at the nose, suggesting that HSN transmits fertility status to the nervous system. HSN also modulates other downstream circuits, such as the nose touch sensory neurons ASH and FLP, which are required for evasion of male contact at the nose. Taken together, we identified a signaling cascade wherein presence vs. absence of sperm is transmitted via monoaminergic signaling from the somatic gonad to the nervous system. This demonstrates how changes in internal state, such as fertility status, act to modulate neural circuits and alter the valence of sensory input.

## Introduction

It is essential for animals to successfully reproduce. While in most cases this requires the participation of two individuals, some animals have evolved mechanisms wherein a self-fertile individual can give rise to progeny independently of a mating partner. In the hermaphroditic nematode *C. elegans*, the hermaphrodite is a somatically female animal that produces sperm for a restricted period of time during larval development, and then switches to oocyte production in adulthood^1,2^. These sperm are stored in the spermatheca and used to fertilize the developing oocytes^3^. While the number of sperm are rate-limiting for the number of progeny that the hermaphrodite can produce, male animals can also mate with a hermaphrodite and contribute additional sperm^4^. Thus, the number of offspring can be greatly increased by successful mating with a male.

However, in *C. elegans*, mating early in adulthood has well-documented risks for the hermaphrodite, including accelerated aging and death^5,6^. Thus, it is initially beneficial for hermaphrodites to avoid male mating attempts. Indeed, it has been shown that self-fertile hermaphrodites move more quickly and are less likely to be mated than sperm-exhausted hermaphrodites^7–9^. This is at least partially due to increased attractiveness of sperm-depleted hermaphrodites to male mating partners^8^. Concurrently, it has been suggested that a behavioral change contributes, as sperm-depleted hermaphrodites are also less likely to flee male mating attempts^7^.

More generally, this indicates that fertility status is an internal state in *C. elegans* that has broad effects on animal physiology and behavior. Internal states in reproductive behavior are the motivation or arousal states that influence how an individual responds to potential mates^10,11^. They can be controlled by a variety of mechanisms, including changes in gene expression^12^, recurrent dynamics^13^, and neuromodulation^14^. The fact that there is a behavioral difference between self-fertile and sperm-depleted *C. elegans* hermaphrodites suggests that the transition in fertility status alters the nervous system.

We sought to determine what initial stimulus from a male mating attempt was aversive to a self-fertile hermaphrodite, and how declining fertility dampens the response to this stimulus. We found that self-fertile *C. elegans* hermaphrodites perform escape behaviors in response to two distinct touch stimuli that occur during the course of male mating: contact of the male tail with the hermaphrodite nose, and the vulva. This escape response depends on monoaminergic signaling from the somatic gonad, which transmits the sperm status to the rest of the nervous system via neuropeptidergic signaling in the HSN neuron.

## Results

To understand the responses of self-fertile and sperm-depleted hermaphrodites to mating attempts, we observed hermaphrodite behavior during canonical mating assays. *C. elegans* males perform a stereotyped series of steps to mate a hermaphrodite animal^15^(**Figure 1A**). When assessing the responses of self-fertile hermaphrodites during these stereotyped steps of male behavior, hermaphrodites would often exhibit escape responses at two particular steps: male turns around the nose, and male contact at the vulva. To quantify this behavior, we split hermaphrodite escape into these two categories (nose evasion and vulva evasion), and calculated an evasion index for each type of response. We found that self-fertile (day 1) hermaphrodites were indeed more likely to evade both nose and vulva contact from males than sperm-exhausted (day 5) hermaphrodites(**Figure 1B, Supplemental Videos 1 and 2**).

**Figure 1:**
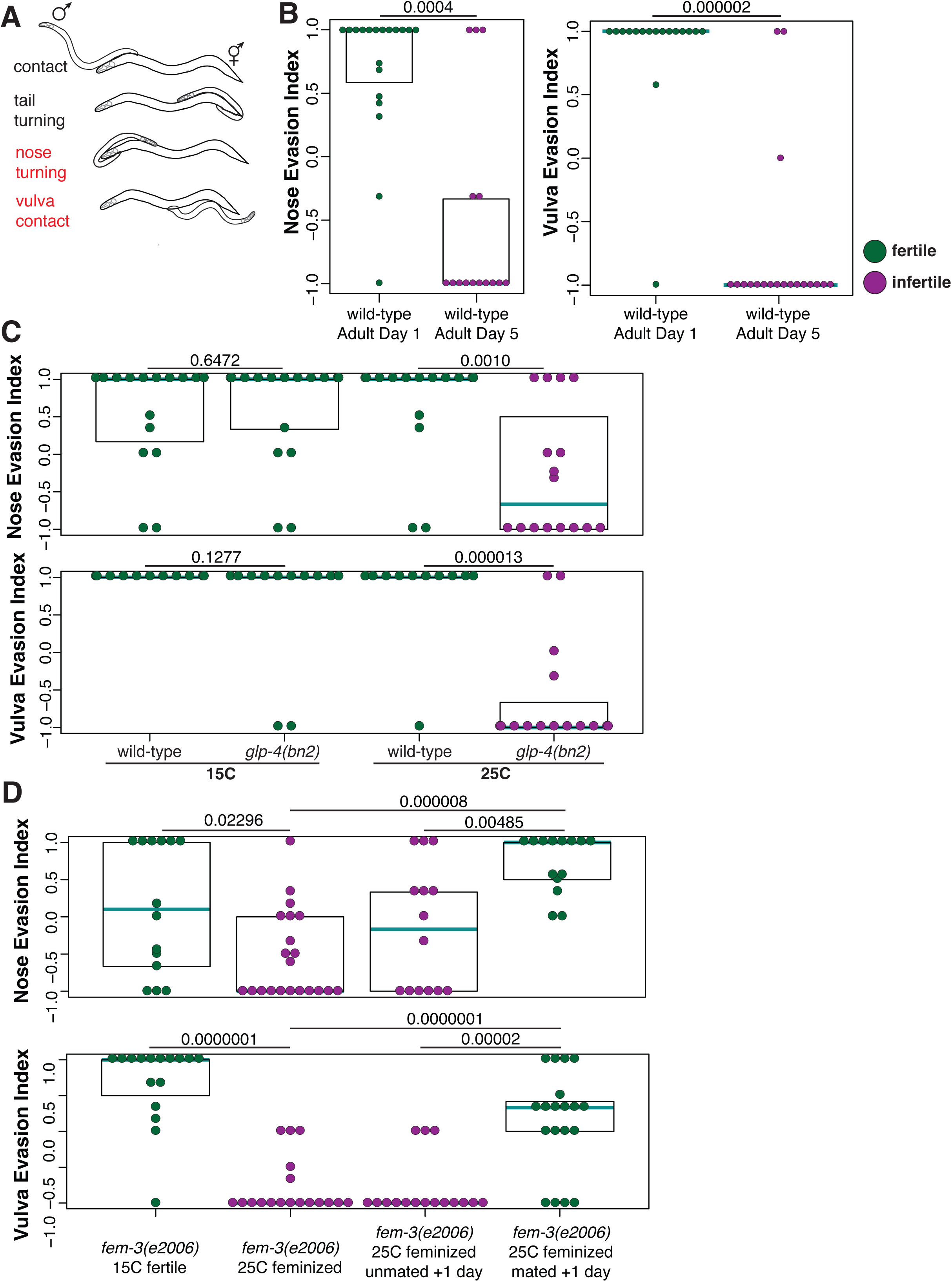
Presence of sperm induces *C. elegans* hermaphrodites to evade mating attempts at the nose and vulva. **A:** Schematic of *C. elegans* mating. Males contact the hermaphrodite with their tail, and then systematically scan the body to search for the vulva. Fertile hermaphrodites escape from turning around the nose or contact at the vulva (red), but not other steps of the process. **B:** Hermaphrodites that have depleted their self-sperm are responsive to mating. We calculated hermaphrodite evasiveness to contact at the nose and vulva individually (see Methods). Dots are color-coded by fertility status of the hermaphrodite, green for fertile and purple for infertile. In all panels, each dot shows the index from one hermaphrodite, cyan lines show medians, black boxes show quartiles. p-values shown above plot calculated by Wilcoxon rank sum test. **C:** Young adult hermaphrodites without a germline are already receptive to mating. All hermaphrodites are day 1 adults. The *glp-4(bn2)* mutation is temperature-sensitive^17^, and mutants have a germline when raised at 15C, but not at 25C. Wild-type control hermaphrodites were also raised at the corresponding temperature (15C or 25C). **D:** Mating evasion is sperm-dependent and reversible. The *fem-3(e2006)* mutation is termperature-sensitive^18^, and hermaphrodites are feminized at 25C but not at 15C. *fem-3(e2006)* 15C fertile and *fem-3(e2006)* 25C feminized hermaphrodites are both day 1 adults, *fem-3(e2006)* 25C feminized unmated +1 day and *fem-3(e2006)* 25C feminized mated +1 day are both day 2 adults.

While day 5 hermaphrodites generally do not yet show signs of systemic decline^16^, it remained possible that this reduction in behavioral evasion was an indirect effect of some aging-related process rather than a specific change related to fertility. We first eliminated the germline completely using the temperature-sensitive *glp-4(bn2)*, wherein germ cells never proliferate due to a defect in mitosis^17^. In the absence of germ cells, we found that day 1 hermaphrodites were already receptive to male mating attempts, suggesting that the germline or fertility is necessary (**Figure 1C**). However, complete removal of the germline impacts many distinct cell types (mitotic germ cells, oocytes, sperm), in addition to physical forces in the uterus caused by the presence of eggs. To disambiguate these effects, we first tested removal of sperm, but not oocytes. Using *fem-3(e2006)* mutants, which are germline-feminized in a temperature-sensitive manner^18^, we found that day 1 germline-feminized hermaphrodites were receptive to male contact at the nose and vulva(**Figure 1D**). This also enabled us to test the plasticity of this behavior. We found that the day after a *fem-3(e2006)* hermaphrodite was mated (ie a fertile day 2 adult hermaphrodite), her behavior reversed and she began to exhibit escape responses from additional male mating attempts, whereas a day 2 *fem-3(e2006)* hermaphrodite that was still a virgin remained receptive to male mating(**Figure 1D**). Thus, we conclude that sperm are necessary for hermaphrodite evasion of male mating, and that this is a reversible behavioral state. We also tested *fem-3(q20)* mutants, which is a temperature-sensitive germline masculinizing allele^19^, and found that these hermaphrodites were constitutively evasive of male mating, even though they were not actually fertile (**Supplemental Figure 1A**). This suggests that the presence of sperm is also sufficient for hermaphrodite evasion of male mating, and that this behavior does not require actual fertility or egg-laying.

Having established that hermaphrodite evasion behavior is modulated by sperm status, we sought to determine how this difference is transmitted to the nervous system. In *C. elegans,* the germline is contained within somatic gonad tissue^3^. The somatic gonad is made up of multiple different cell types, which both form the anatomical structure of the gonad and control the development and maintenance of the germline^20–22^. In *gon-2(q388)* mutants (which are temperature-sensitive), the gonad progenitors fail to divide and differentiate at the restrictive temperature of 25C, which also prevents germline differentiation^23^. Thus, we expected that these gonadless mutants should become receptive to male mating attempts as day 1 hermaphrodites. Interestingly, we found that even though these mutants do not have sperm and should not evade male mating attempts, they remained fully evasive to male contact at the vulva, and partially evasive to male contact at the nose(**Figure 2A**). One interpretation of this result is that some signal from the somatic gonad normally acts as a repressor of hermaphrodite evasion in the absence of sperm, and that without the somatic gonad this behavior cannot be repressed.

**Figure 2:**
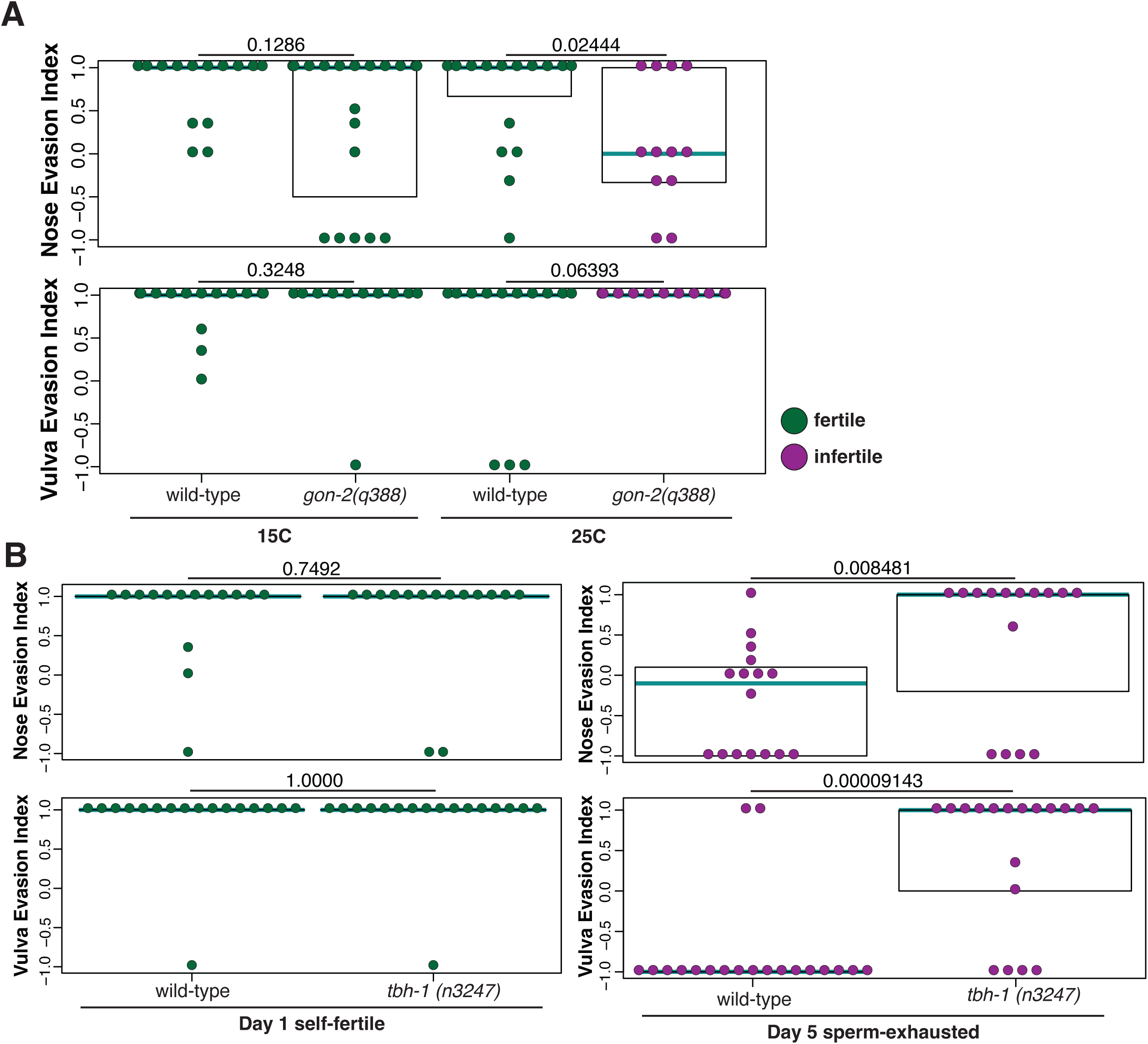
Octopaminergic signaling from the gonadal sheath suppresses evasion. **A:** Gonadless hermaphrodites evade mating attempts even though they are infertile. *gon-2(q388)* hermaphrodites have a gonad when raised at 15C (left) but are gonadless when raised at 25C (right). Wild-type control hermaphrodites were also raised at the corresponding temperature (15C or 25C). Dots are color-coded by fertility status of the hermaphrodite, green for fertile and purple for infertile. In all panels, each dot shows the index from one hermaphrodite, cyan lines show medians, black boxes show quartiles. p-values shown above plot calculated by Wilcoxon rank sum test. There is a small decrease in evasiveness to nose contact in gonadless hermaphrodites, suggesting that an additional parallel pathway may influence hermaphrodite response at the nose. **B:** Hermaphrodites lacking octopamine production evade mating attempts even though they are infertile. *tbh-1(n3247)* mutant hermaphrodites lack the enzyme required to convert tyramine to octopamine^28^. Self-fertile hermaphrodites have normal mating evasion (left), whereas sperm-exhausted hermaphrodites continue to evade male mating attempts (right).

Similar to our finding that the presence of sperm triggers hermaphrodite avoidance behavior, the somatic gonad also autonomously responds to the presence versus absence of sperm to control oocyte maturation^24^. This function in controlling oocyte maturation depends on octopamine, which is produced by the gonad sheath cells^25^. Octopamine, a monoamine neurotransmitter which is considered the invertebrate analog of norepinephrine^26^, also has wide-ranging roles in the *C. elegans* nervous system^27^. Thus, we assessed hermaphrodites defective in the synthesis of octopamine (mutants for the *tdc-1* or *tbh-1* enzymes). We found that loss of either of these enzymes did not affect self-fertile hermaphrodites: they still showed normal evasion from male contact at both the nose and vulva(**Figure 2B, Supplemental Figure 1B**). However, sperm-exhausted hermaphrodites were unable to suppress this behavior without the production of octopamine, and continued to inappropriately evade male mating attempts(**Figure 2B**). This inappropriate evasion was very robust for vulva contact, and more variable for nose contact, similar with our intermediate results in the *gon-2(q388)* mutant. The gonad sheath is one of two places that octopamine is generated in the worm, the other being the RIC interneuron^28^. To disambiguate octopamine from these two sources, we rescued *tbh-1* in RIC, and found that these hermaphrodites were still inappropriately evasive(**Supplemental Figure 1C**).

Having identified the germline and gonad components required for mating evasion, we wondered how the evasion behavior is modulated by the nervous system. Vulva innervation in *C. elegans* has been well-studied in the context of the egg-laying circuit. Egg-laying is regulated by two neuron classes, the HSN and VC neurons, in addition to neuroendocrine cells^29^. The HSN neurons can be genetically ablated using the *egl-1(n986)* mutation, which causes them to undergo inappropriate apoptosis during embryogenesis^30^. In *egl-1* mutants, we found that self-fertile hermaphrodites were not only receptive to male contact at the vulva, but also at the nose (**Figure 3A**). In contrast, silencing the VC neurons had no effect on hermaphrodite escape behavior (**Figure 3B**). HSN has a cell body and local innervation at the vulva, and then projects directly into the nerve ring (**Figure 3C**). This poises it both to respond locally to gonad signals but also transmit this information broadly to the nervous system.

**Figure 3:**
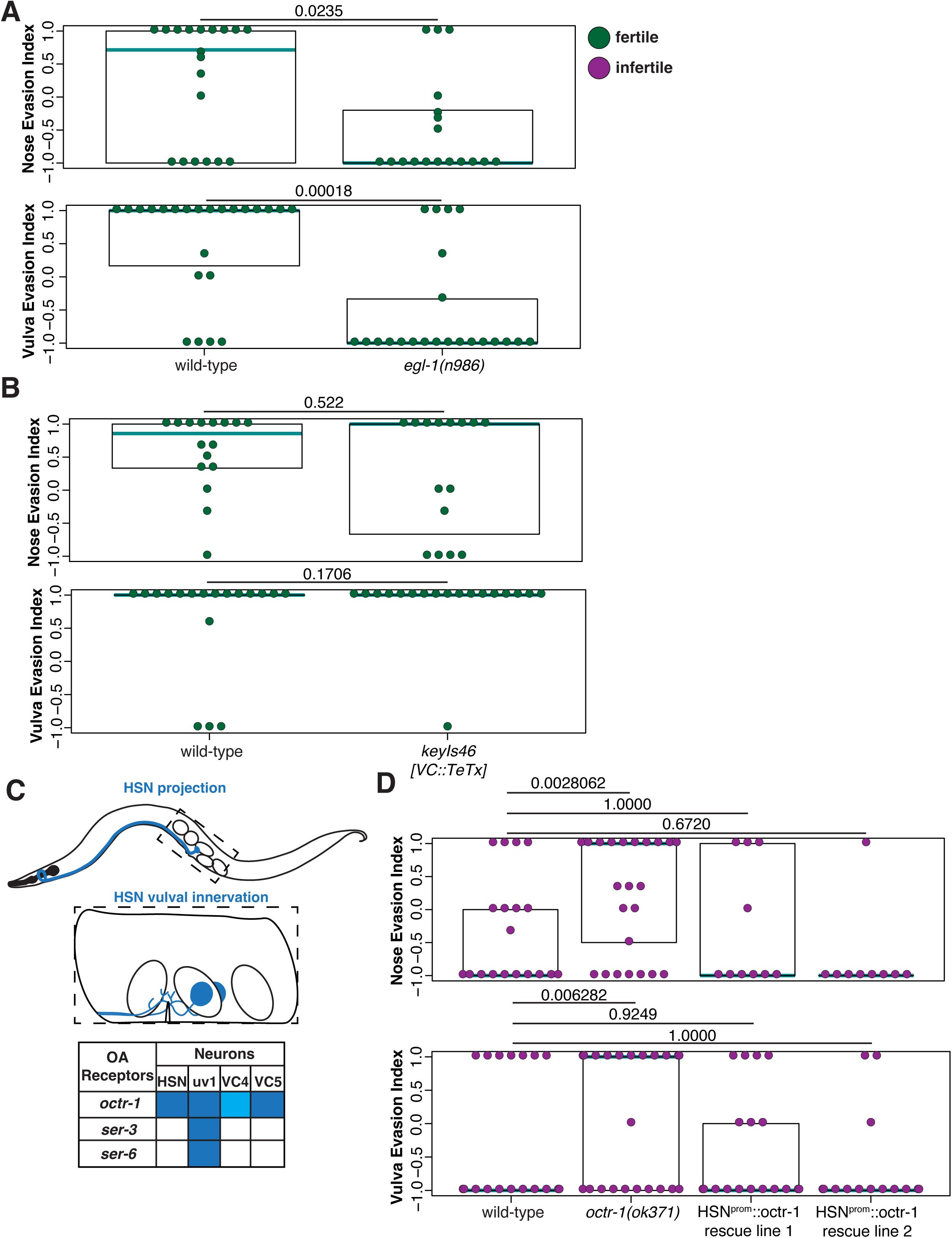
HSN responds to fertility status via octopamine signaling. **A:** Genetic ablation of HSN causes inappropriate receptivity to mating. The *egl-1(n986)* mutation causes HSN to undergo apoptosis during embryonic development^30^. Dots are color-coded by fertility status of the hermaphrodite, green for fertile and purple for infertile. In all panels, each dot shows the index from one hermaphrodite, cyan lines show medians, black boxes show quartiles. p-values shown above plot calculated by Wilcoxon rank sum test. **B:** VC neurons are not required for male evasion behavior. Day 1 self-fertile hermaphrodites with transgenically silenced VC neurons^45^ are still evasive of male mating attempts. **C:** Schematics of HSN axon projection into the nerve ring (top), branching innervation at the vulva (middle) and octopamine receptor expression in the egg-laying circuitry (bottom). HSN schematics are adapted from wormatlas.org, table summarizing octopamine receptor expression is adapted from^31^. **D:** *octr-1* is required in HSN for receptivity of hermaphrodites to male mating attempts. In day 5 sperm-depleted hermaphrodites, *octr-1(ok371)* mutants maintain significant evasiveness to mating attempts, which is rescued by expressing *octr-1* using an HSN-specific *cat-4* promoter^56^. p-values shown above plot calculated by Wilcoxon rank sum test followed by Bonferroni correction for multiple testing.

Because of its proximity to the vulva, we wondered whether HSN cell-autonomously responds to octopamine from the gonad sheath. *C. elegans* has 3 known octopamine receptors, and their expression patterns in the egg-laying circuitry have been described^31^. HSN has been shown to express the *octr-1* receptor, suggesting that it could respond directly to the octopamine cue from the somatic gonad^31^(**Figure 3C**). Indeed, we found that *octr-1* mutants phenocopied the loss of octopamine: hermaphrodite behavior was normal while the animal was self-fertile but animals were unable to suppress escape behaviors upon sperm depletion (**Figure 3D**). HSN-specific rescue of *octr-1* restored receptivity of sperm-depleted hermaphrodites to male mating attempts (**Figure 3D**).

To determine whether the gonad also communicates with the nervous system independently of HSN, we combined loss of HSN with loss of the gonad sheath to perform an epistasis experiment. If all signals from the gonad are transmitted via HSN, then we expect that loss of HSN combined with loss of the gonad would result in receptivity to mating. If the gonad also signals via parallel pathways, loss of both HSN and the gonad would maintain evasiveness. We found that that these animals were receptive to male mating attempts (**Supplemental Figure 2A**), suggesting that HSN is the bottleneck for communicating fertility status to the nervous system.

While HSN is well-situated to transmit information about fertility to the nervous system, it does not project to the nose, and thus cannot cell-autonomously respond to male mating contact at the nose. Neurons responsible for response to touch stimuli at the nose have been characterized in *C. elegans*^32^. In the standard nose touch response assay, an eyelash pick is placed in front of a worm moving forward, so that the animal collides perpendicularly with the eyelash. Escape response to these collisions is primarily mediated via the polymodal ASH sensory neuron^32^. We found that when we transgenically silenced ASH, self-fertile hermaphrodites were inappropriately receptive to male contact at the nose, although they still evaded male contact at the vulva(**Figure 4A**). This is in contrast to our genetic ablation of HSN, which impacted both behaviors. This suggests that nose responsiveness is the downstream behavior, and does not feed back onto vulva responsiveness.

**Figure 4:**
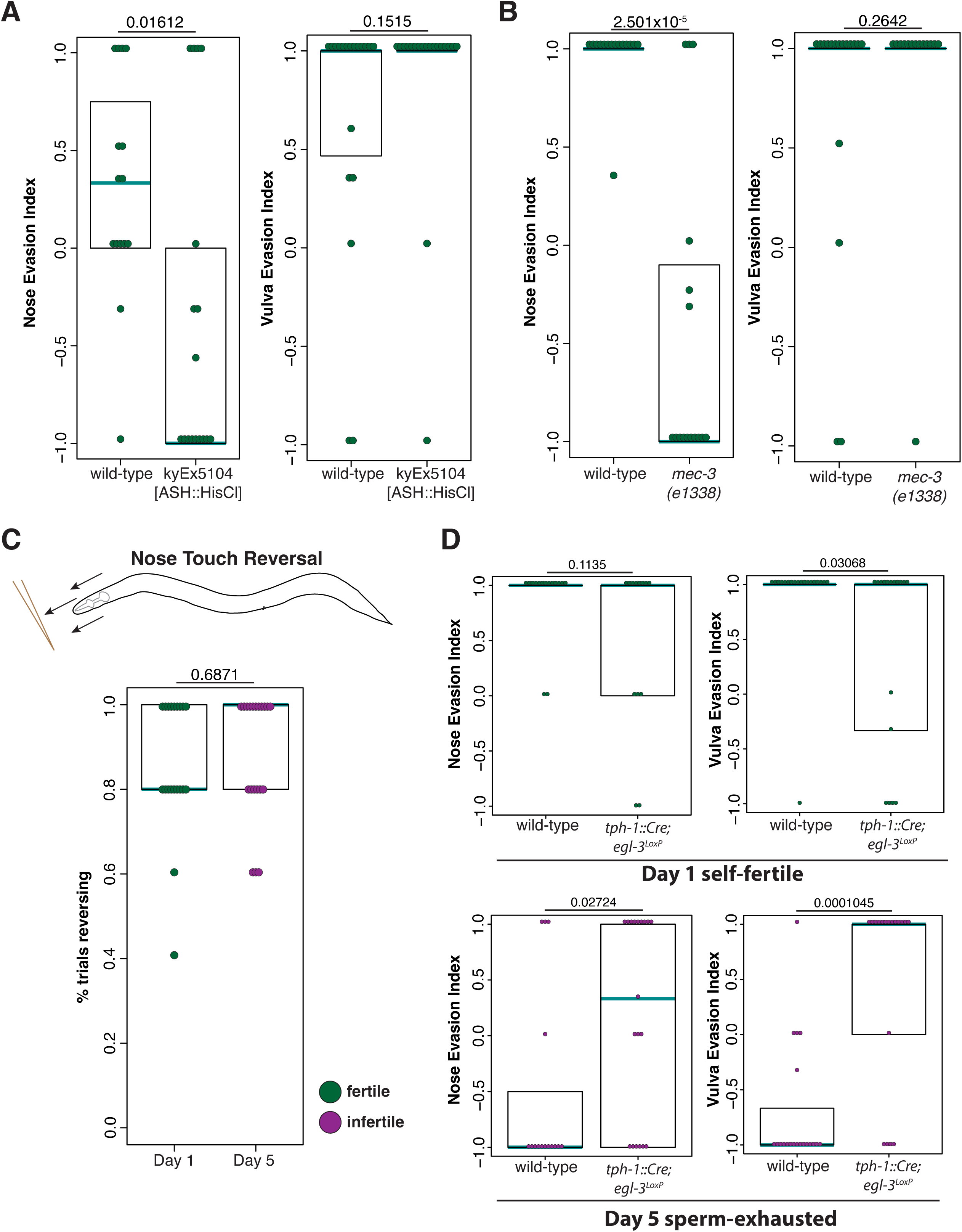
Evasion of male contact at the nose is a behavioral modality modulated by fertility via HSN. A: Silencing the ASH neurons makes hermaphrodites receptive to male mating contact at the nose, but not the vulva. Dots are color-coded by fertility status of the hermaphrodite, green for fertile and purple for infertile. In all panels, each dot shows the index from one hermaphrodite, cyan lines show medians, black boxes show quartiles. p-values shown above plot calculated by Wilcoxon rank sum test. B: Mutants lacking the FLP, PVD, and touch receptor neurons are constitutively receptive to male contact at the nose, but not the vulva. C: Nose touch response is not affected by fertility status. Schematic above of nose touch response assay (collision with an eyebrow while animal is moving forward). Below, each dot represents the performance of a single hermaphrodite in a nose touch response assay. Each hermaphrodite is assayed 5 times (see Methods), and the number of responses is used to generate a percentage of responses. p-value above calculated using Wilcoxon rank sum test. D: Neuropeptide production from HSN is required for mating receptivity in sperm-exhausted hermaphrodites. Self-fertile hermaphrodites (above) evade mating attempts, but do not become receptive to mating when sperm-exhausted (below). There is a small decrease in evasiveness to nose contact in the absence of HSN neuropeptides, suggesting that an additional parallel pathway may influence hermaphrodite response at the nose.

Animals with dominant *mec-4* mutations are defective in touch response to the body due to degeneration of the touch responsive neurons^33^. Loss of these neurons does not affect standard nose touch response^32^, and we similarly found that it did not have an effect on hermaphrodite mating evasion(**Supplemental Figure 2B**). However, hermaphrodites were also inappropriately receptive to male contact at the nose in mutants for *mec-3*, which affects the FLP and PVD sensory neurons in addition to the touch responsive neurons(**Figure 4B**). While the dendritic arbor of PVD covers the body, FLP innervates the head, and was shown as a secondary neuron for detecting standard nose touch^32^. We conclude that both the ASH and FLP neurons mediate response to male contact at the nose in fertile hermaphrodites. Interestingly, we found that sperm-depleted hermaphrodites still had normal response to the standard nose touch assay(**Figure 4C**). This suggests that hermaphrodite response to male contact at the nose is regulated differently than standard nose touch.

Because loss of HSN affected both nose and vulva contact behaviors, whereas loss/silencing of ASH and FLP only affected nose touch behaviors, we reasoned that HSN must be the upstream neuron and signaling to the head sensory neurons. HSN is both cholinergic and serotonergic^34^ and also displays an unusually high degree of neuropeptidergic connectivity^35^. We found that mutants for the serotonin-synthesizing *tph-1* pathway displayed normal male evasion behavior (**Supplemental Figure 2C**).

However, using an intersectional strategy to remove the KPC2 pro-protein convertase *egl-3* in HSN^36^, we found that loss of neuropeptidergic signaling phenocopied loss of octopamine signaling: hermaphrodites without HSN neuropeptides exhibited normal male evasion while self-fertile, but were inappropriately still evasive once sperm-depleted(**Figure 4D**). Consistent with our model that HSN transmits information about fertility status from the germline (via the somatic gonad) to downstream neurons, this manipulation affected both vulva and nose evasion behaviors.

## Discussion

We found that the decreased likelihood of self-fertile hermaphrodites to be mated by males involves a behavioral state, wherein fertile hermaphrodites evade contact by males at the nose and vulva during mating attempts. The presence or absence of sperm is transmitted by the somatic gonad via octopamine to HSN, which plays a central role in modulating hermaphrodite response to male mating attempts. Across species, it has been found that small numbers of neurons can drive reproductive (aggression and courtship) behavioral states^10,11^. For instance, in male *Drosophila*, subsets of the P1 neurons can drive both male courtship behavior^37^ and male-male aggression^38^. Similarly, Esr1+ ventromedial hypothalamic neurons in mouse can drive both male mounting and aggression behavior^39^. Here we report that in *C. elegans* mating, HSN serves as both a link between the gonad to nervous system, and a bottleneck to modulate hermaphrodite mating evasion.

HSN has been previously implicated in a variety of behaviors, such as egg-laying^40,41^, and modulation of locomotion^42,43^. While HSN-ablated animals are still capable of laying eggs, the periodicity of egg-laying is compromised, and this is phenocopied by mutants without serotonin^44^. Optogenetic activation of HSN causes overall locomotory speeding independent of fertility, suggesting that output from HSN can drive general locomotory circuitry^42^. Locally at the vulva, HSN behaves as both a motor and interneuron: neuromuscular junctions onto vulval muscle are required for egg-laying^42^, and HSN can also respond to reciprocal mechanosensory feedback from vulval muscle^45^. HSN is also inhibited by high osmolarity via the BAG neurons^42,43^ and gentle touch via the PLM neuron^43^, suggesting that in some contexts it acts downstream of other sensory neurons. We propose that in controlling hermaphrodite evasion of male mating attempts, HSN is acting as an interneuron between primary sensory inputs at the nose and vulva (from FLP and ASH, and possibly vulval muscle pressure) and upstream of the locomotory circuits to ultimately drive the escape behavior.

One open question is the identity of the signal that the somatic gonad receives from sperm cells. In species that compete for mating opportunities, males can utilize chemical cues to suppress further mating by females. In both *D. melanogaster* and *A. aegypti*, males transfer sex peptides during mating that alter females’ behavior to prevent remating^46,47^. The best-studied protein released from *C. elegans* sperm is the filamentous protein known as Major Sperm Protein (MSP), which is encoded by a family of dozens of genes and controls oocyte maturation and gonad sheath contraction^48,49^.

While it is difficult to genetically eliminate MSP due to the redundancy among the large gene family, it has been shown that hermaphrodites mutant for the *vab-1* Ephrin receptor have defects in oocyte maturation and sheath contraction^50^. Specifically, while oocyte maturation and sheath contraction are not properly repressed once sperm are exhausted, they still occur when sperm are present. However, feminized *fog-2(q71)* hermaphrodites, which lack sperm entirely, also do not mature oocytes or have gonad sheath contractions^24^. Thus, in addition to MSP there is an initial positive signal from sperm that induces oocyte maturation and sheath contraction. While study of *C. elegans* fertility has revealed a variety of conserved sperm proteins, it is believed that there are still many sperm peptides yet unidentified^51^. It is possible that these fertility-relevant sperm proteins also contribute to hermaphrodite mating evasiveness. In contrast to systems like *D. melanogaster* and *A. aegypti,* where mating suppression is conferred as a competitive tactic by male animals, we found that both self- and cross-sperm suppress mating in *C. elegans*. This enables *C. elegans* hermaphrodites to extend their reproductive span when necessary, but avoid deleterious health and lifespan effects of mating early in adulthood.

Hermaphroditism in *C. elegans* evolved from an ancestral gonochoristic (male-female) system, and this occurred multiple times independently in *Caenorhabditis*^52^. While hermaphroditic species can evade male mating attempts without sacrificing reproduction, this is not true of the gonochoristic ancestors. Surprisingly, study of gonochoristic species in the *Caenorhabditis* genus has suggested that attempted escape by females from male mating attempts may be the evolutionarily default state, but these males have evolved the ability to paralyze females during mating^53^. This paralytic ability by males, and the hermaphroditic sensitivity to it, has been lost in *C. elegans*^53^. While the initial evasiveness of *C. elegans* hermaphrodites could be evolutionarily advantageous in light of the health risks of early mating^5,6^, it is less apparent why evasion would exist as the default state in male-female species. It has been shown that some inter-specific hybridizations are possible within the quickly-evolving *Caenorhabditis* genus, but that these hybridizations do not give rise to successful populations^54^. Default evasion of mates by *Caenorhabditis* females could thus serve to protect from non-specific mating, and provide another checkpoint for species specificity with the evolution of male paralysis and female receptivity.

The nervous system can alter its function to allow flexibility in response to internal states as diverse as fear, hunger, or social isolation^11^. Fertility is one such internal state, which can impact how animals interact with each other (aggression, courtship) and the environment (nutritional needs, nest-building)^10,55^. Here we show that in *C. elegans*, fertility modulates how hermaphrodites respond to tactile contact from males. This state persists as long as sperm are present, and can be reversed by mating. We were able to define a signaling pathway that links sperm status, via the somatic gonad, to the HSN neuron, and then to sensory and escape circuitry in the rest of the nervous system.

## Acknowledgements

We thank members of the Hobert and Bayer labs for feedback on the manuscript. Strains were generously provided by the Hobert, Bargmann, and Collins labs, and by the CGC, which is supported by the NIH Office of Research Infrastructure Programs (P40 OD010440). This research was supported by the University of Basel and New York University. EAB was a fellow of the Jane Coffin Childs Memorial Fund for Medical Research.

## Materials Availability

Novel strains will be deposited at the Caenorhabditis Genetics Center or are available from the lead contact upon request.

## Author Contributions

Conceptualization, Investigation, Funding Acquisition, and Writing – Original Draft: EAB. Investigation and Writing – Review and Editing: IV.

## Data and Code Availability

Data supporting the findings of this study are available within the paper and its Supplementary Information. Detailed descriptions of novel strains generated in this study are given in the Methods section. This paper does not report original code.

## Methods

### C. elegans husbandry

Worms were maintained by standard methods^2^. All strains were maintained at 20C with the exception of temperature-sensitive strains (detailed below). Wild-type hermaphrodites (as indicated in figures) are genotype N2 or *him-5(e1490)*. We found that the mating avoidance and nose touch response behaviors of day 1 and day 5 N2 and *him-5(e1490)* hermaphrodites are indistinguishable. To generate day 1 adult hermaphrodites for behavioral assays, we picked L4 hermaphrodites that had been raised at 20C and allowed them to acclimate overnight at assay (room) temperature. Day 5 hermaphrodites were maintained at 20C and transferred each day away from the brood until day 4 of adulthood, when they were acclimated overnight at assay (room) temperature.

The following alleles and transgenes were used in this study:

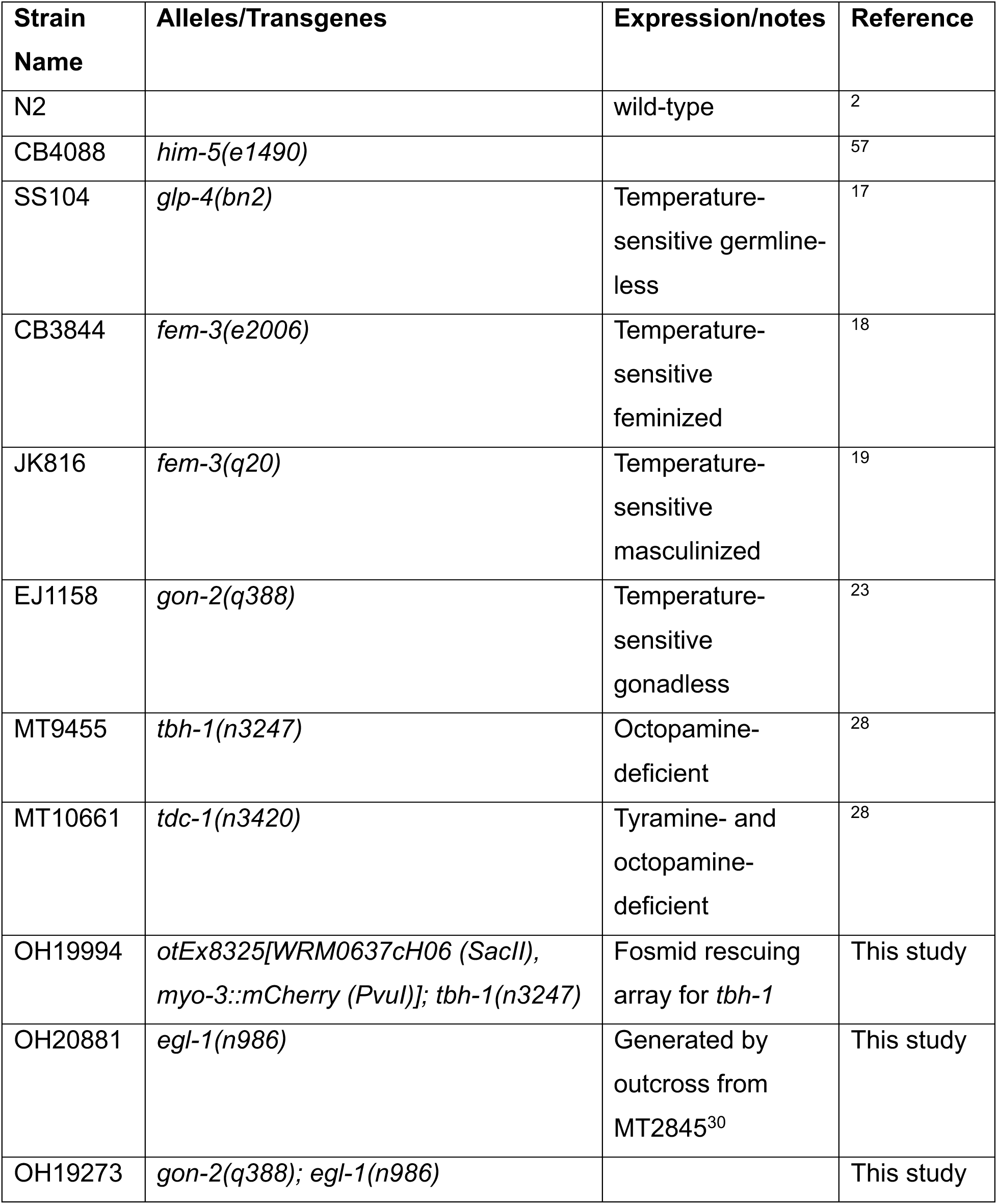
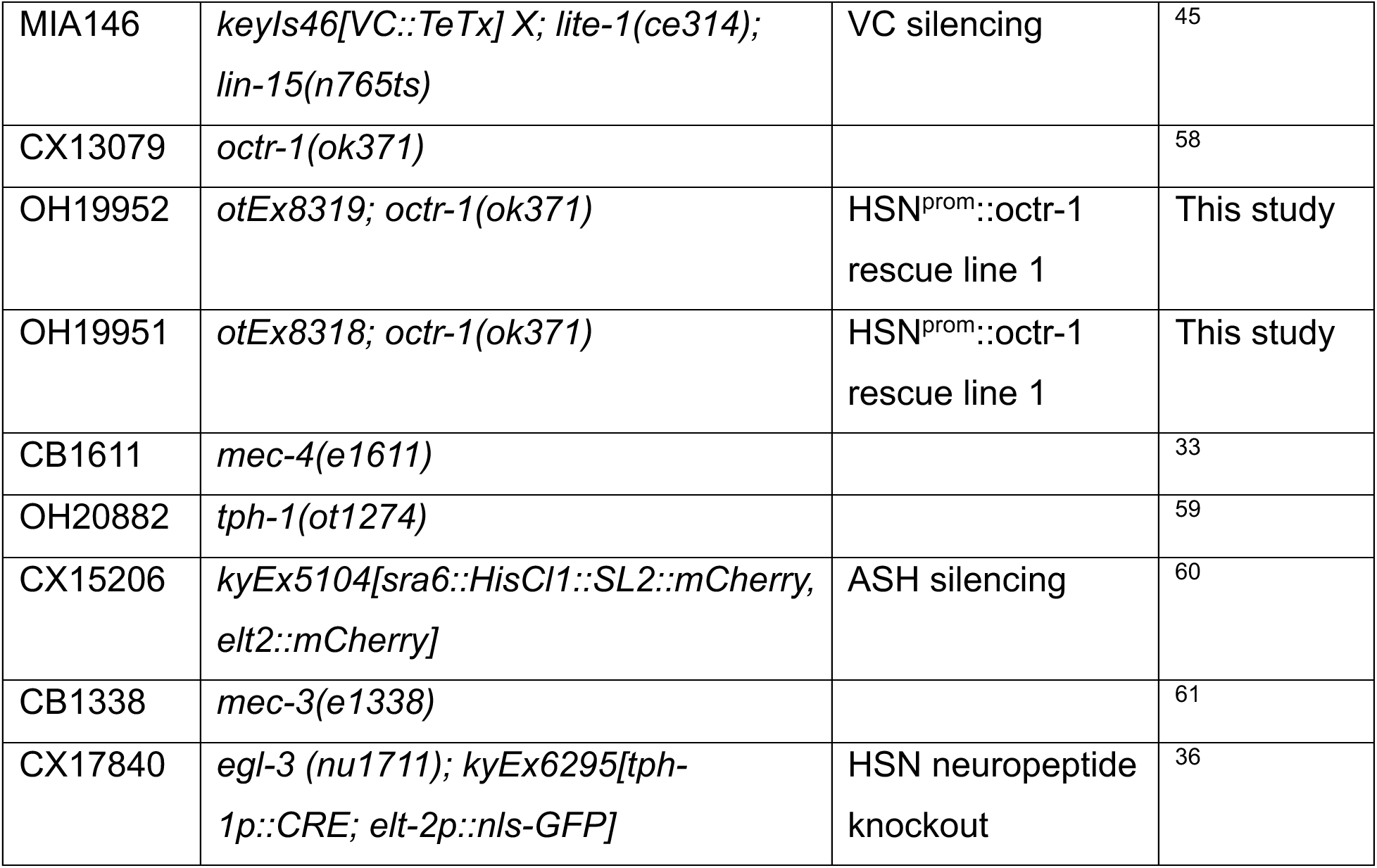

### Temperature-sensitive strains

*glp-4(bn2)* germline-less*, fem-3(e2006)* feminized, and *fem-3(q20)* masculinized mutants are all temperature-sensitive. Phenotypic adults were generated by picking gravid mutant adults (raised at 15C) to 25C, and progeny were manually selected from the following generation for assay. For the *fem-3(e2006)* assay, where we also scored behavior in day 2 adults that had been mated, group mating plates were set up with day 1 adults for one hour and then hermaphrodites were separated from males. The following day, we sorted fertile hermaphrodites (mated) from hermaphrodites that were still infertile (unmated) for assay.

*gon-2(q388)* gonad-less adults were generated by picking gravid mutant adults (raised at 15C) to 25C, and manually selecting progeny from the following generation for assay. We found that this temperature shift caused a range of phenotypic severity in the progeny, likely due to the known maternal contribution of *gon-2* mRNA or protein^23^. Complete loss of *gon-2* function results in animals that are vulvaless in addition to gonadless^23^, which would preclude mating behavior. We thus selected animals that had a vulva but had defective gonad development: these animals were always sterile, and had a small “blob” of gonad tissue in the midbody but no discernible gonad differentiation or structure.

### Mating evasion assay

Mating plates were prepared with a small bacterial lawn (10uL drop of OP50) 1-2 days in advance. 10-15 day 1 adult him-5 males are transferred to the plate, then hermaphrodites are assayed one at a time. The assay lasts until sperm transfer occurs, or until 10 minutes of failed mating attempts, at which point the hermaphrodite is removed and the male that successfully copulated is replaced, if there was one. The evasion index is calculated similarly to *C. elegans* chemotaxis indices: (evasions - nonevasions)/total touches. For instance, if a male contacted the vulva 5 times, and the hermaphrodite escaped from the first 4 attempts but successful mating occurred on the 5th contact, the evasion score would be (4-1)/5=0.6. Head contact and vulva contact are counted separately, and the evasion index is calculated for each individually.

Hermaphrodite evasion from nose contact is always a reversal, and evasion from vulva contact is often a reversal, but sometimes the hermaphrodite simply sprints forward without reversing first (or will sprint forward and then reverse to change direction later). Regardless of the directionality, all evasive maneuvers were scored as an evasion. Often the male entirely loses contact with the hermaphrodite during these evasions, but this was not necessary for a maneuver to be considered evasive.

10 hermaphrodites were assayed per replicate, and each experiment is the combination of at least two replicates (on separate days). However, some animals will have null values (ie a male contacted the hermaphrodite directly on the ventral side and successfully mated with the vulva without performing a turn around the nose, so there would be a 0/0 for the nose evasion index). For some experiments involving comparing several genotypes in parallel (such as rescue experiments), replicates were performed across multiple days. Control animals were always included with every parallel replicate. All experiments were performed blinded to hermaphrodite genotype.

### Histamine-gated chloride channel silencing

ASH::HisCl1 (kyEx5104) transgenic animals were picked at the L4 stage and placed on NGM plates containing 10 mM histamine with OP50 bacteria as a food source. Wild-type animals were also placed on 10mM histamine plates at L4, to control for any effects of the histamine on behavior. Mating assays were performed as detailed above, but also on 10mM histamine plates.

### Nose touch response assay

Nose touch response assays were performed according to^32^. An eyelash pick was placed in front of an animal that was moving forward, and an animal was considered responding if it reversed following this collision. Each animal was trialed 5 times, with at least 5 minutes between each trial.

## Reference

1 Nigon, V. Les modalités de la reproduction et le déterminisme du sexe chez quelques Nématodes libres. Ann Sci Nat Zool 11, 1–132 (1949).

2 Brenner, S. The genetics of Caenorhabditis elegans. Genetics 77, 71–94 (1974). 10.1093/genetics/77.1.71

3 Hirsh, D., Oppenheim, D. & Klass, M. Development of the reproductive system of Caenorhabditis elegans. Dev Biol 49, 200–219 (1976). 10.1016/0012-1606(76)90267-0

4 Hodgkin, J. Male Phenotypes and Mating Efficiency in CAENORHABDITIS ELEGANS. Genetics 103, 43–64 (1983). 10.1093/genetics/103.1.43

5 Shi, C. & Murphy, C. T. Mating induces shrinking and death in Caenorhabditis mothers. Science 343, 536–540 (2014). 10.1126/science.1242958

6 Gems, D. & Riddle, D. L. Longevity in Caenorhabditis elegans reduced by mating but not gamete production. Nature 379, 723–725 (1996). 10.1038/379723a0

7 Kleemann, G. A. & Basolo, A. L. Facultative decrease in mating resistance in hermaphroditic Caenorhabditis elegans with self-sperm depletion. Animal Behaviour 74, 1339–1347 (2007). 10.1016/j.anbehav.2007.02.031

8 Morsci, N. S., Haas, L. A. & Barr, M. M. Sperm status regulates sexual attraction in Caenorhabditis elegans. Genetics 189, 1341–1346 (2011). 10.1534/genetics.111.133603

9 Suo, S. Sperm function is required for suppressing locomotor activity of C. elegans hermaphrodites. PLoS One 19, e0297802 (2024). 10.1371/journal.pone.0297802

10 Anderson, D. J. Circuit modules linking internal states and social behaviour in flies and mice. Nat Rev Neurosci 17, 692–704 (2016). 10.1038/nrn.2016.125

11 Flavell, S. W., Gogolla, N., Lovett-Barron, M. & Zelikowsky, M. The emergence and influence of internal states. Neuron 110, 2545–2570 (2022). 10.1016/j.neuron.2022.04.030

12 Zhang, S. X., Rogulja, D. & Crickmore, M. A. Recurrent Circuitry Sustains Drosophila Courtship Drive While Priming Itself for Satiety. Curr Biol 29, 3216–3228 e3219 (2019). 10.1016/j.cub.2019.08.015

13 Deutsch, D., et al. The neural basis for a persistent internal state in Drosophila females. Elife 9 (2020). 10.7554/eLife.59502

14 Tayler, T. D., Pacheco, D. A., Hergarden, A. C., Murthy, M. & Anderson, D. J. A neuropeptide circuit that coordinates sperm transfer and copulation duration in Drosophila. Proc Natl Acad Sci U S A 109, 20697–20702 (2012). 10.1073/pnas.1218246109

15 Liu, K. S. & Sternberg, P. W. Sensory regulation of male mating behavior in Caenorhabditis elegans. Neuron 14, 79–89 (1995). 10.1016/0896-6273(95)90242-2

16 Herndon, L. A., et al. Stochastic and genetic factors influence tissue-specific decline in ageing C. elegans. Nature 419, 808–814 (2002). 10.1038/nature01135

17 Beanan, M. J. & Strome, S. Characterization of a germ-line proliferation mutation in C. elegans. Development 116, 755–766 (1992). 10.1242/dev.116.3.755

18 Hodgkin, J. Sex determination in the nematode C. elegans: analysis of tra-3 suppressors and characterization of fem genes. Genetics 114, 15–52 (1986). 10.1093/genetics/114.1.15

19 Barton, M. K., Schedl, T. B. & Kimble, J. Gain-of-function mutations of fem-3, a sex-determination gene in Caenorhabditis elegans. Genetics 115, 107–119 (1987). 10.1093/genetics/115.1.107

20 Kimble, J. E. & White, J. G. On the control of germ cell development in Caenorhabditis elegans. Dev Biol 81, 208–219 (1981). 10.1016/0012-1606(81)90284-0

21 Hall, D. H., et al. Ultrastructural features of the adult hermaphrodite gonad of Caenorhabditis elegans: relations between the germ line and soma. Dev Biol 212, 101–123 (1999). 10.1006/dbio.1999.9356

22 McCarter, J., Bartlett, B., Dang, T. & Schedl, T. Soma-germ cell interactions in Caenorhabditis elegans: multiple events of hermaphrodite germline development require the somatic sheath and spermathecal lineages. Dev Biol 181, 121–143 (1997). 10.1006/dbio.1996.8429

23 Sun, A. Y. & Lambie, E. J. gon-2, a gene required for gonadogenesis in Caenorhabditis elegans. Genetics 147, 1077–1089 (1997). 10.1093/genetics/147.3.1077

24 McCarter, J., Bartlett, B., Dang, T. & Schedl, T. On the control of oocyte meiotic maturation and ovulation in Caenorhabditis elegans. Dev Biol 205, 111–128 (1999). 10.1006/dbio.1998.9109

25 Kim, J., Hyun, M., Hibi, M. & You, Y. J. Maintenance of quiescent oocytes by noradrenergic signals. Nat Commun 12, 6925 (2021). 10.1038/s41467-021-26945-x

26 Roeder, T. Octopamine in invertebrates. Prog Neurobiol 59, 533–561 (1999). 10.1016/s0301-0082(99)00016-7

27 Horvitz, H. R., Chalfie, M., Trent, C., Sulston, J. E. & Evans, P. D. Serotonin and octopamine in the nematode Caenorhabditis elegans. Science 216, 1012–1014 (1982). 10.1126/science.6805073

28 Alkema, M. J., Hunter-Ensor, M., Ringstad, N. & Horvitz, H. R. Tyramine Functions independently of octopamine in the Caenorhabditis elegans nervous system. Neuron 46, 247–260 (2005). 10.1016/j.neuron.2005.02.024

29 Collins, K. M., et al. Activity of the C. elegans egg-laying behavior circuit is controlled by competing activation and feedback inhibition. Elife 5 (2016). 10.7554/eLife.21126

30 Desai, C. & Horvitz, H. R. Caenorhabditis elegans mutants defective in the functioning of the motor neurons responsible for egg laying. Genetics 121, 703–721 (1989). 10.1093/genetics/121.4.703

31 Fernandez, R. W., et al. Cellular Expression and Functional Roles of All 26 Neurotransmitter GPCRs in the C. elegans Egg-Laying Circuit. J Neurosci 40, 7475–7488 (2020). 10.1523/JNEUROSCI.1357-20.2020

32 Kaplan, J. M. & Horvitz, H. R. A dual mechanosensory and chemosensory neuron in Caenorhabditis elegans. Proc Natl Acad Sci U S A 90, 2227–2231 (1993). 10.1073/pnas.90.6.2227

33 Driscoll, M. & Chalfie, M. The mec-4 gene is a member of a family of Caenorhabditis elegans genes that can mutate to induce neuronal degeneration. Nature 349, 588–593 (1991). 10.1038/349588a0

34 Weinshenker, D., Garriga, G. & Thomas, J. H. Genetic and pharmacological analysis of neurotransmitters controlling egg laying in C. elegans. J Neurosci 15, 6975–6985 (1995). 10.1523/JNEUROSCI.15-10-06975.1995

35 Ripoll-Sanchez, L., et al. The neuropeptidergic connectome of C. elegans. Neuron 111, 3570–3589 e3575 (2023). 10.1016/j.neuron.2023.09.043

36 Marquina-Solis, J., et al. Antagonism between neuropeptides and monoamines in a distributed circuit for pathogen avoidance. Cell Rep 43, 114042 (2024). 10.1016/j.celrep.2024.114042

37 Kimura, K., Hachiya, T., Koganezawa, M., Tazawa, T. & Yamamoto, D. Fruitless and doublesex coordinate to generate male-specific neurons that can initiate courtship. Neuron 59, 759–769 (2008). 10.1016/j.neuron.2008.06.007

38 Hoopfer, E. D., Jung, Y., Inagaki, H. K., Rubin, G. M. & Anderson, D. J. P1 interneurons promote a persistent internal state that enhances inter-male aggression in Drosophila. Elife 4 (2015). 10.7554/eLife.11346

39 Lee, H., et al. Scalable control of mounting and attack by Esr1+ neurons in the ventromedial hypothalamus. Nature 509, 627–632 (2014). 10.1038/nature13169

40 Trent, C., Tsung, N. & Horvitz, H. R. Egg-laying defective mutants of the nematode Caenorhabditis elegans. Genetics 104, 619–647 (1983). 10.1093/genetics/104.4.619

41 Brewer, J. C., Olson, A. C., Collins, K. M. & Koelle, M. R. Serotonin and neuropeptides are both released by the HSN command neuron to initiate Caenorhabditis elegans egg laying. PLoS Genet 15, e1007896 (2019). 10.1371/journal.pgen.1007896

42 Huang, Y. C., et al. A single neuron in C. elegans orchestrates multiple motor outputs through parallel modes of transmission. Curr Biol 33, 4430–4445 e4436 (2023). 10.1016/j.cub.2023.08.088

43 Zhang, M., et al. A self-regulating feed-forward circuit controlling C. elegans egg-laying behavior. Curr Biol 18, 1445–1455 (2008). 10.1016/j.cub.2008.08.047

44 Waggoner, L. E., Zhou, G. T., Schafer, R. W. & Schafer, W. R. Control of alternative behavioral states by serotonin in Caenorhabditis elegans. Neuron 21, 203–214 (1998). 10.1016/s0896-6273(00)80527-9

45 Medrano, E. & Collins, K. M. Muscle-directed mechanosensory feedback activates egg-laying circuit activity and behavior in Caenorhabditis elegans. Curr Biol 33, 2330–2339 e2338 (2023). 10.1016/j.cub.2023.05.008

46 Yapici, N., Kim, Y. J., Ribeiro, C. & Dickson, B. J. A receptor that mediates the post-mating switch in Drosophila reproductive behaviour. Nature 451, 33–37 (2008). 10.1038/nature06483

47 Duvall, L. B., Basrur, N. S., Molina, H., McMeniman, C. J. & Vosshall, L. B. A Peptide Signaling System that Rapidly Enforces Paternity in the Aedes aegypti Mosquito. Curr Biol 27, 3734–3742 e3735 (2017). 10.1016/j.cub.2017.10.074

48 Ward, S., et al. Genomic organization of major sperm protein genes and pseudogenes in the nematode Caenorhabditis elegans. J Mol Biol 199, 1–13 (1988). 10.1016/0022-2836(88)90374-9

49 Miller, M. A., et al. A sperm cytoskeletal protein that signals oocyte meiotic maturation and ovulation. Science 291, 2144–2147 (2001). 10.1126/science.1057586

50 Miller, M. A., Ruest, P. J., Kosinski, M., Hanks, S. K. & Greenstein, D. An Eph receptor sperm-sensing control mechanism for oocyte meiotic maturation in Caenorhabditis elegans. Genes Dev 17, 187–200 (2003). 10.1101/gad.1028303

51 Mei, X. & Singson, A. W. The molecular underpinnings of fertility: Genetic approaches in Caenorhabditis elegans. Adv Genet (Hoboken) 2 (2021). 10.1002/ggn2.10034

52 Kiontke, K., et al. Caenorhabditis phylogeny predicts convergence of hermaphroditism and extensive intron loss. Proc Natl Acad Sci U S A 101, 9003–9008 (2004). 10.1073/pnas.0403094101

53 Garcia, L. R., LeBoeuf, B. & Koo, P. Diversity in mating behavior of hermaphroditic and male-female Caenorhabditis nematodes. Genetics 175, 1761–1771 (2007). 10.1534/genetics.106.068304

54 Devi, M. P., et al. Five new Caenorhabditis species from Indonesia provide exceptions to Haldane’s rule and partial fertility of interspecific hybrids. G3 (Bethesda) 15 (2025). 10.1093/g3journal/jkaf134

55 Tinbergen, N. On the analysis of social organization among vertebrates, with special reference to birds. (University of Notre Dame P, 1939).

56 Lloret-Fernandez, C., et al. A transcription factor collective defines the HSN serotonergic neuron regulatory landscape. Elife 7 (2018). 10.7554/eLife.32785

57 Hodgkin, J., Horvitz, H. R. & Brenner, S. Nondisjunction Mutants of the Nematode CAENORHABDITIS ELEGANS. Genetics 91, 67–94 (1979). 10.1093/genetics/91.1.67

58 Flavell, S. W., et al. Serotonin and the neuropeptide PDF initiate and extend opposing behavioral states in C. elegans. Cell 154, 1023–1035 (2013). 10.1016/j.cell.2013.08.001

59 Liao, C. P., Majeed, M. & Hobert, O. Experience-dependent, sexually dimorphic synaptic connectivity defined by sex-specific cadherin expression. Sci Adv 10, eadq9183 (2024). 10.1126/sciadv.adq9183

60 Oren-Suissa, M., Bayer, E. A. & Hobert, O. Sex-specific pruning of neuronal synapses in Caenorhabditis elegans. Nature 533, 206–211 (2016). 10.1038/nature17977

61 Xue, D., Tu, Y. & Chalfie, M. Cooperative interactions between the Caenorhabditis elegans homeoproteins UNC-86 and MEC-3. Science 261, 1324–1328 (1993). 10.1126/science.8103239

