## Supplementary material for "Fertility reversibly modulates *C. elegans* behavior via gonad-nervous system signaling": Document S1

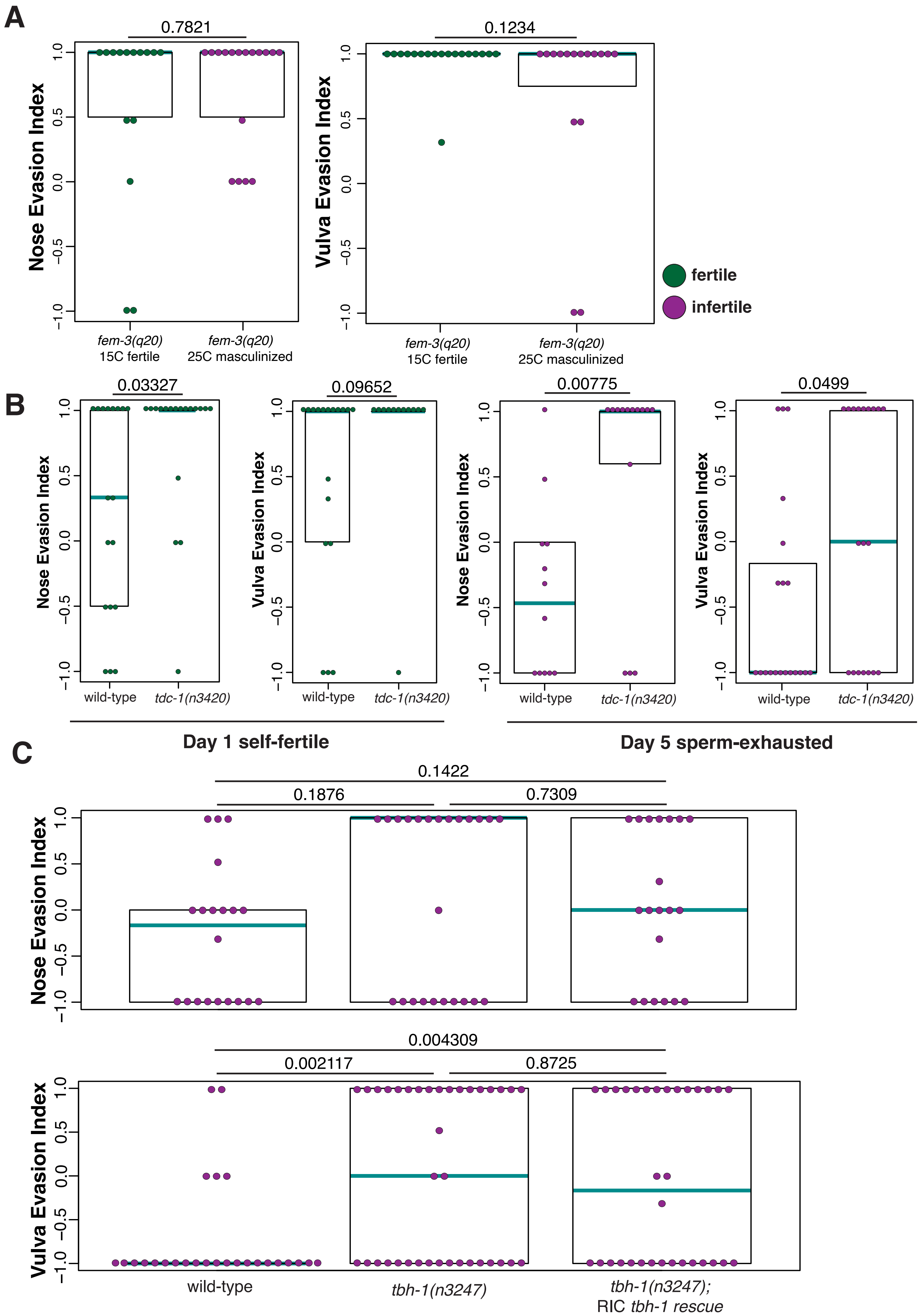

A
0.7821
Nose Evasion Index
fem-3(q20)
15C fertile
fem-3(q20)
25C masculinized
0.1234
Vulva Evasion Index
fem-3(q20)
15C fertile
fem-3(q20)
25C masculinized
fertile
infertile
B
0.03327
Nose Evasion Index
wild-type
tdc-1(n3420)
0.09652
Vulva Evasion Index
wild-type
tdc-1(n3420)
0.00775
Nose Evasion Index
wild-type
tdc-1(n3420)
0.0499
Vulva Evasion Index
wild-type
tdc-1(n3420)
Day 1 self-fertile
Day 5 sperm-exhausted
C
0.1422
0.1876
0.7309
Nose Evasion Index
0.004309
0.002117
0.8725
Vulva Evasion Index
wild-type
tbh-1(n3247)
tbh-1(n3247);
RIC tbh-1 rescue

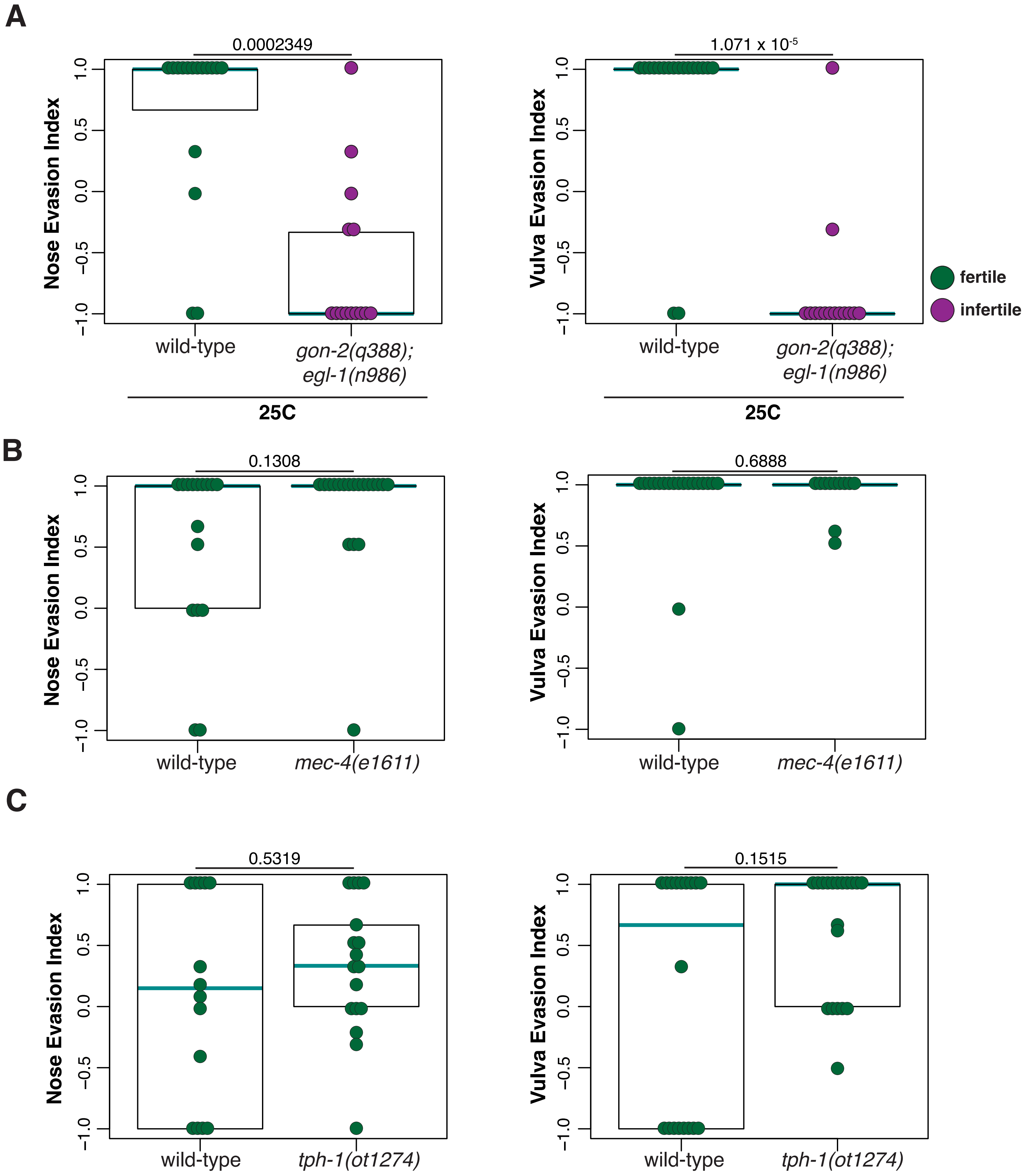

A
0.0002349
Nose Evasion Index
wild-type
gon-2(q388); egl-1(n986)
25C
1.071 x 10-5
Vulva Evasion Index
fertile
infertile
B
0.1308
mec-4(e1611)
0.6888
C
0.5319
tph-1(ot1274)
0.1515

**Supplemental Figure 1: Sperm and octopamine convey the fertility signal.**

**A:** Germline masculinized hermaphrodites are evasive of male mating attempts. *fem-3(q20)* hermaphrodites are fertile when raised at 15C but produce only sperm when raised at 25C. Dots are color-coded by fertility status of the hermaphrodite, green for fertile and purple for infertile. In all panels, each dot shows the index from one hermaphrodite, cyan lines show medians, black boxes show quartiles. p-values shown above plot calculated by Wilcoxon rank sum test.

**B:** *tdc-1(n3420)* mutants are evasive of male mating attempts even when sperm-exhausted. *tdc-1(n3420)* mutants lack the tyrosine decarboxylase enzyme and cannot generate tyramine or octopamine[1]. Self-fertile hermaphrodites (left) evade mating attempts, but do not become receptive to mating when sperm-exhausted (right).

**C:** Expression of *tbh-1* in the RIC neurons does not rescue evasiveness in sperm-depleted hermaphrodites. In contrast to **Figure 2B,** *tbh-1* mutants showed a strongly bimodal phenotype in these replicates. Vulva evasion of *tbh-1* mutants was still significant by Wilcoxon rank sum test in these replicates, but nose evasion was not. However, due to the increased bimodality in the *tbh-1* mutants, the distribution between wild-type and *tbh-1* mutants remained significant for nose evasion (p=0.0273 by Kolmogorov-Smirnov test).

**Supplemental Figure 2: HSN serves as a bottleneck for mating evasion, which does not require the touch receptor neurons or serotonin.**

**A:** *gon-2; egl-1* double mutants are receptive to male mating attempts, suggesting that HSN acts downstream of the somatic gonad in regulating hermaphrodite mating evasion. In all panels, each dot shows the index from one hermaphrodite, cyan lines show medians, black boxes show quartiles. p-values shown above plot calculated by Wilcoxon rank sum test. All animals were day 1 hermaphrodites raised at 25C (see Methods).

**B:** *mec-4(e1611)* mutants have defective touch receptor neurons[2], but normal mating evasion behavior. All animals were day 1 hermaphrodites.

**C:** *tph-1(ot1274)* mutants do not produce serotonin, and have normal mating evasion behavior. All animals were day 1 hermaphrodites.

**Supplemental Video 1: Fertile hermaphrodites evade male mating attempts.** Video of a day 1 adult *him-5(e1490)* male attempting to mate a day 1 wild-type (N2) hermaphrodite. The hermaphrodite performs reversals in response to two male turns around the nose (32s and 41s), and then sprints away from contact at the vulva (47s) to ultimately escape the mating attempt.

**Supplemental Video 2: Sperm-exhausted hermaphrodites are receptive to mating.** Video of a day 1 adult *him-5(e1490)* male mating a day 5 adult wild-type (N2) hermaphrodite. The hermaphrodite does not evade from a male turn around the nose (30s) or contact at the vulva (37s).

**Reference:**


1 Alkema, M. J., Hunter-Ensor, M., Ringstad, N. & Horvitz, H. R. Tyramine Functions independently of octopamine in the Caenorhabditis elegans nervous system. *Neuron* **46**, 247–260 (2005). https://doi.org/10.1016/j.neuron.2005.02.024

2 Driscoll, M. & Chalfie, M. The mec-4 gene is a member of a family of Caenorhabditis elegans genes that can mutate to induce neuronal degeneration. *Nature* **349**, 588–593 (1991). https://doi.org/10.1038/349588a0
